# Integrated Expression Analysis of C-MYC Oncogene-Associated Pathways in Gastric Adenocarcinoma and it’s Correlation with Clinicopathological Factors

**DOI:** 10.1101/2025.09.11.675674

**Authors:** Kauê Sant’Ana Pereira Guimarães, Murilo Filho Pereira Marinho, Jennifer Somtochukwu Nwoye, Ronald Matheus da Silva Mourão, Juliana Barreto Albuquerque Pinto, Jéssica Manoelli Costa da Silva, Diego Pereira, Valéria Cristiane Santos da Silva, Samia Demachki, Williams Fernandes Barra, Samir Mansour Casseb, Fabiano Cordeiro Moreira, Rommel Mario Rodriguez Burbano, Paulo Pimentel de Assumpção

## Abstract

**Background:** The C-MYC oncogene is a well-established driver of gastric carcinogenesis, yet the integrated expression pattern of its complex regulatory network and its clinical implications in gastric adenocarcinoma (GAC) remain to be fully elucidated. This study aimed to perform an integrated bioinformatic analysis of C-MYC and its associated pathways in a cohort of GAC patients to delineate its expression profile, assess its potential as a biomarker, and correlate its patterns with clinicopathological factors such as Laurén classification and neoadjuvant treatment status.

**Methods:** The study included transcriptome data from 74 GAC tumor samples and 68 normal gastric tissue samples. Following RNA sequencing, a comprehensive bioinformatic analysis was conducted on a curated list of 21 C-MYC-associated genes. The methodology included differential expression analysis (DESeq2), unsupervised hierarchical clustering, Principal Component Analysis (PCA), and Receiver Operating Characteristic (ROC) curve analysis to evaluate diagnostic performance. Gene expression levels were also statistically correlated with Laurén histological subtypes and neoadjuvant therapy status using the Wilcoxon rank-sum test.

**Results:** The analysis revealed a profound dysregulation of the C-MYC network in GAC. While *MYC* itself was significantly upregulated, its transcriptional antagonists, particularly *MXD4* and *MXD3*, were the most significantly downregulated genes. This gene signature robustly separated tumor from normal tissues in both hierarchical clustering and PCA. ROC analysis demonstrated the outstanding diagnostic potential of several genes, with *MXD4* achieving a perfect Area Under the Curve (AUC) of 1.00, surpassing the diagnostic value of *MYC* (AUC=0.86). Stratification by Laurén classification showed that *MYC* and its stability regulator *PTBP1* were significantly more expressed in the intestinal subtype, whereas the repressors *MXD3* and *MXI1* were higher in the diffuse subtype. No significant expression differences were observed based on neoadjuvant treatment status.

**Conclusion:** Gastric adenocarcinoma is characterized by a coordinated dysregulation of the C-MYC network, marked by both oncogene activation and a concurrent loss of its key transcriptional repressors. The profound downregulation of antagonists like *MXD4* serves as an exceptionally accurate molecular signature for GAC, suggesting its potential as a diagnostic biomarker superior to *MYC* alone. The divergent expression patterns between Laurén subtypes highlight distinct molecular pathobiology and may have implications for targeted therapies

## Introduction

Gastric carcinogenesis is recognized as a multifactorial process in which genetic and environmental factors interact, activating multiple intracellular signaling pathways that culminate in uncontrolled cell growth (1). Central to many of these alterations is the dysregulation of the proto-oncogene C-MYC, one of the most crucial oncogenes in human cancer, which is found to be dysregulated in over 50% of tumors, including gastric adenocarcinoma (2,3).

The C-MYC gene regulates multiple biological processes, primarily acting as a transcription factor that, through dimerization with the MAX protein (MYC Associated Factor X), binds to specific DNA sequences known as E-boxes in the promoter region of target genes (CACGFG) to modulate gene expression. Upon binding, the MYC-MAX complex recruits other proteins including Histone acetyltransferases (HATs), and TIP43 - an ATP-binding protein to open up chromatin and increase the expression of target genes. This protein complex activates genes critical for growth, proliferation, and the cell cycle, while suppressing the transcription of genes that inhibit mitotic processes, such as CDKN1A (p21) (4,5).

The activity of C-MYC is finely regulated by a network of antagonistic interactions with the MAD family of proteins. During cellular differentiation, a molecular switch happens where the MYC protein is downregulated and displaced by the MAD family, forming MAD-MAX dimers. By binding to MAX to form MAD-MAX dimers, it functions as transcriptional repressors, silencing genes activated by MYC and allowing cells from various tissues to enter a post-mitotic, differentiated state (6). However, genetic deregulation of MYC expression and the loss of checkpoint components, such as tumor suppressor TP53, prevents this control, allowing MYC to drive malignancy (7).

In addition to this central axis, the oncogenic function of C-MYC is amplified by complementary pathways. Recent studies demonstrate that C-MYC directly interacts with the protein arginine methyltransferase 5 (PRMT5) to transcriptionally repress a cohort of tumor suppressor genes, including PTEN and CDKN1A, thereby promoting the growth of gastric cancer cells (8). Its protein stability, in turn, is controlled by an axis involving the PTBP1 protein, which prevents C-MYC degradation mediated by the ubiquitin ligase FBW7. This mechanism is crucial for maintaining cancer stem cell (CSC) phenotypes, which are responsible for chemoresistance and relapse (9). Furthermore, C-MYC orchestrates immune evasion by regulating the expression of checkpoint ligands, such as CD274 (PD-L1) and CD47, and modulating cytokines to create an immunosuppressive tumor microenvironment (2).

Despite the knowledge of these individual pathways, the collective expression pattern of these different C-MYC-associated regulators and their integrated clinical relevance in gastric adenocarcinoma are not yet fully elucidated. Therefore, the objective of this study is to perform an integrated bioinformatic analysis of the expression of C-MYC, its main partner and target genes in a cohort of gastric cancer patients. We aim to delineate the expression profile of these pathways, evaluate their potential as biomarkers, and correlate their patterns with important clinicopathological features, including Laurén histological classification and neoadjuvant treatment status.

## Materials and Methods

### Ethics statement

This study was approved by the Ethics and Research Committee of João de Barros Barreto University Hospital (approval number: 47580121.9.0000.5634) and was conducted in accordance with the principles outlined in the Declaration of Helsinki. Participant recruitment and sample collection were carried out between July 2, 2022, and July 6, 2023. Before enrollment, all participants received detailed information about the study’s objectives, potential benefits, risks, and possible harms, ensuring a thorough understanding of the research. Written informed consent was voluntarily obtained from all participants prior to their inclusion in the study.

### Sample Collection and Cohort Assembly

Tumor and peritumoral (non-tumoral, adjacent to the tumor at a distance of 5 cm) samples were collected from 72 patients treated at a hospital located in the State of Pará, Brazil. The study was approved by the Research Ethics Committee under protocol number CAAE 47580121.9.0000.5634. Additionally, to expand the cohort of normal tissues, transcriptome data from gastric tissues of cancer-free patients were obtained from the PRJNA1054173 project, publicly available in the NCBI BioProject database. The final cohort for this study consisted of 74 gastric adenocarcinoma (GC) samples and 68 normal gastric tissue samples.

### RNA Sequencing and Data Processing

RNA sequencing (RNA-seq) was performed on the NextSeq® platform using the NextSeq® 500 MID Output V2 kit. In the dry-lab stage, the initial processing of raw sequencing reads included demultiplexing, followed by quality filtering and adapter trimming using Fastp (v0.23.2). Subsequently, reads were aligned and quantified using Salmon against the human transcriptome (Gencode version 43).

### Data Import, Filtering, and Normalization

Gene expression data were imported into R software (v4.3.2) using the tximport package, which converted transcript quantifications into a gene-level count matrix. A filter was applied to retain genes with a minimum of 10 reads in at least 80% of the samples, ensuring that only genes with relevant expression levels were included in the analysis.

Subsequently, the count data were normalized using the rlog (regularized logarithm) function from the DESeq2 package. This approach stabilizes the variance across the mean, making the data suitable for comparative analyses. An additional normalization step was performed using the MinMaxScaler from the scikit-learn library (accessed via the reticulatepackage) to scale the expression data of all samples to a [0, 1] range.

### Bioinformatic Analysis

#### Selection of C-MYC-Associated Genes

Based on a literature review of the mechanisms governing C-MYC’s oncogenic activity, a consolidated list of 21 genes was selected for analysis. This list included core members of the MYC/MAX/MAD axis, key regulators of C-MYC protein stability (e.g., PTBP1, FBXW7), transcriptional partners (PRMT5), and mediators of immune evasion (CD274, CD47).

#### Differential Expression and Visualization

Differential gene expression analysis between GC and normal tissues was performed using the DESeq2 package. Genes were considered significantly differentially expressed if they met the criteria of an absolute log2(Fold Change) > 1 and an adjusted p-value (padj) < 0.05. The results were visualized using a Volcano Plot. An unsupervised hierarchical clustering heatmap, based on the Z-score of rlog-normalized expression values, was created to visualize expression patterns across samples. Principal Component Analysis (PCA) was performed to assess the ability of the gene signature to distinguish between sample groups.

#### Receiver Operating Characteristic (ROC) Analysis

To evaluate the individual diagnostic performance of each gene in the selected signature, ROC curve analysis was performed using the pROC package. The area under the curve (AUC) was calculated for each gene to measure its ability to discriminate between GC and normal tissues.

#### Correlation with Clinicopathological Data

The rlog-normalized expression levels of the selected genes within the tumor cohort were stratified by two clinical variables: Laurén histological classification (Intestinal vs. Diffuse) and neoadjuvant treatment status (Yes vs. No). The statistical significance of expression differences between groups was assessed using the Wilcoxon rank-sum test, and results were visualized using violin plots.

## Results

### Differential Expression of C-MYC-Associated Genes

Differential expression analysis between gastric cancer (GC) and normal tissues revealed a profound and widespread dysregulation of C-MYC-associated genes, as visualized in the Volcano Plot **(Figure 1)**. A central finding was the statistically significant upregulation of the MYC oncogene, which presented a log2(Fold Change) of approximately 1.5. This result, confirming the central role of MYC in GC biology, is consistent with previous studies, such as that of Souza et al. (2013), who reported that 77% of gastric tumors exhibited C-MYC immunoreactivity, which was associated with increased mRNA expression, deeper tumor extension, and the presence of metastasis. Along with MYC, other genes such as FBXW7, a ubiquitin ligase that regulates MYC degradation, and TP63 were also significantly upregulated. Conversely, the analysis demonstrated a robust downregulation of several transcriptional repressors and regulatory partners. The MYC antagonists MXD4 and MXD3 were the most significantly downregulated genes, with a log2(Fold Change) of approximately −2.0 and −1.8, respectively, and extremely high statistical significance. Other key members of this axis, such as MAX, MXI1, and CDKN1C, were also significantly downregulated, indicating that the loss of the repressive machinery is as prominent a feature as the activation of MYC itself.

**Figure 1:**
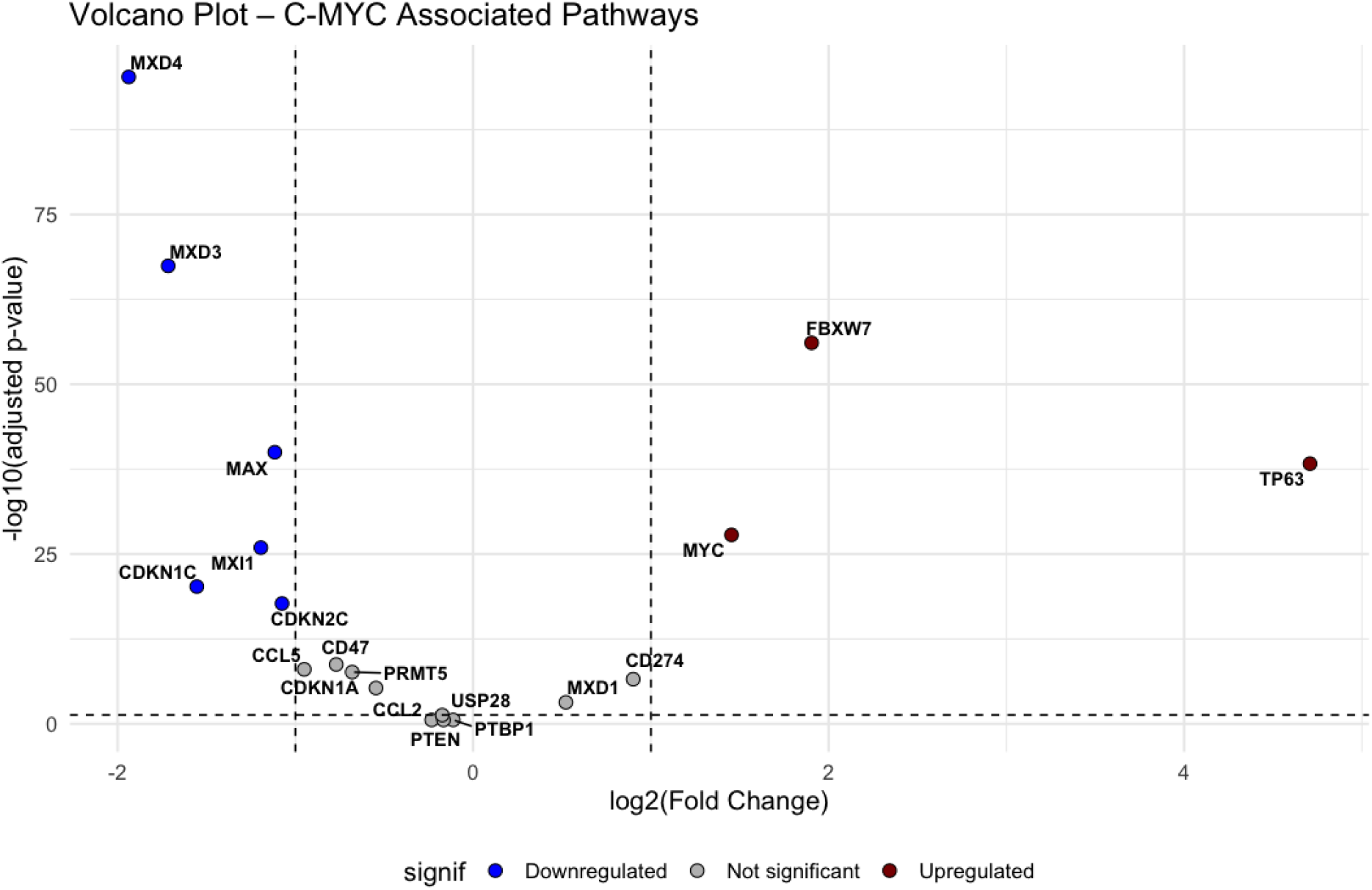
Volcano Plot – Pathways Associated with C-MYC: Volcano plot of differential gene expression between gastric cancer (GC) tissues and normal tissues.

### Gene Expression Patterns Distinguish Tumor from Normal Tissues

To evaluate the collective behavior and expression patterns of this gene signature acros all samples, unsupervised hierarchical clustering was performed, as visualized in a heatmap **(Figure 2)**. The analysis revealed a remarkable segregation of the samples into two distinct primary clusters, which perfectly corresponded to the GC tissue (red) and healthy normal tissue (teal) groups. This unequivocal separation demonstrates that the differential expression of these C-MYC-associated genes constitutes a robust and consistent molecular signature of gastric cancer. Within the heatmap, blocks of genes with coordinated expression patterns can be observed. One gene cluster, including FBXW7, PTEN, and MYC itself, exhibits a predominantly higher expression pattern (red tones) in the GC cohort. In opposition, a larger gene cluster containing the repressors MAX, MXD3, CDKN1A, and PRMT5, shows a predominantly lower expression pattern (blue tones) in the tumor tissues. This coordinated behavior suggests complete reprogramming of this regulatory axis during gastric carcinogenesis.

**Figure 2:**
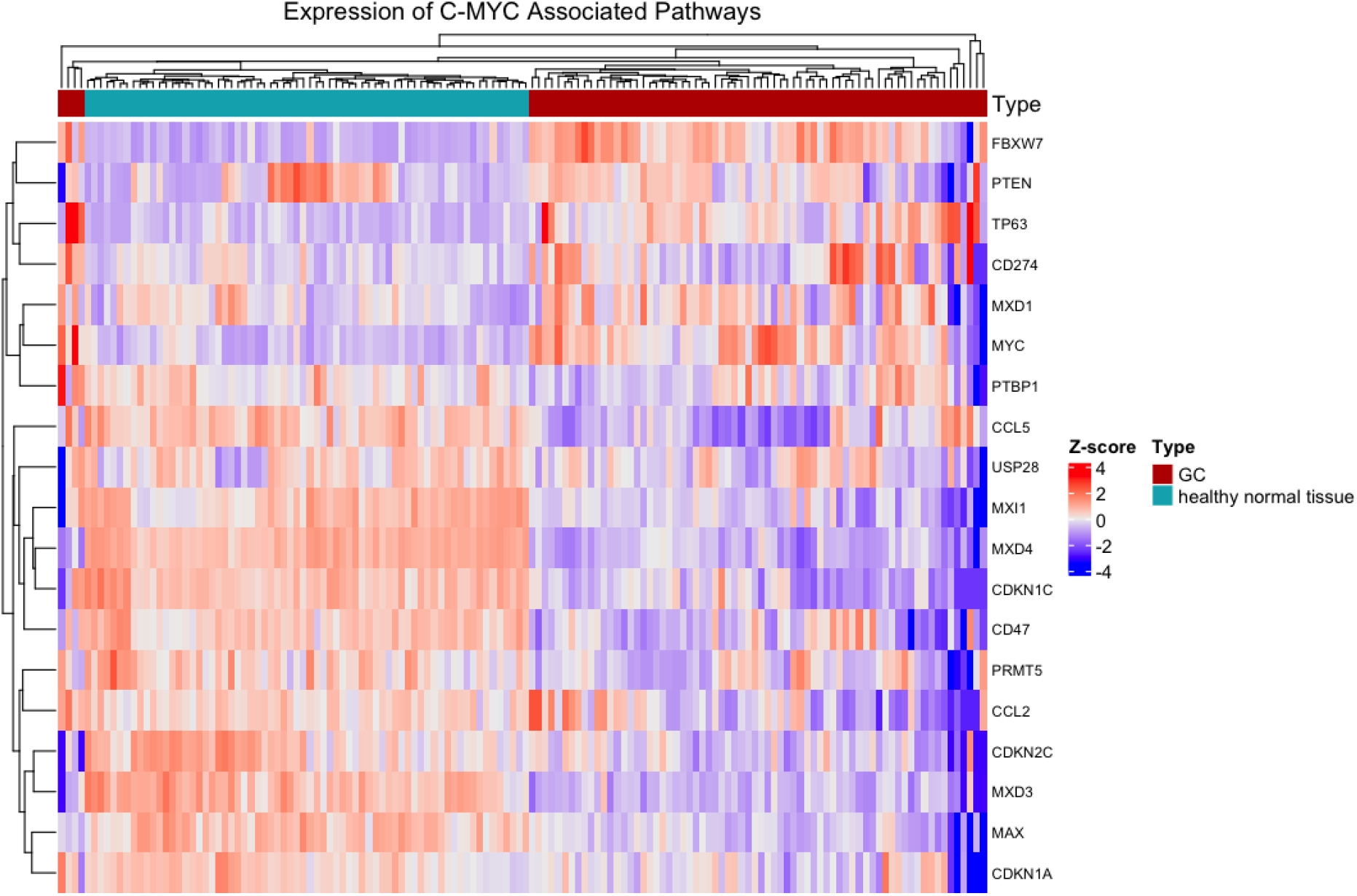
Expression of Pathways Associated with C-MYC: Hierarchical clustering heatmap of the expression of 21 genes associated with C-MYC across all samples.

### Principal Component Analysis (PCA)

Confirming the results of the hierarchical clustering, Principal Component Analysis (PCA) demonstrated the strength of this gene signature in distinguishing the sample types **(Figure 4)**. The PCA plot, which reduces the multidimensional complexity of the expression data into two principal components, showed that the tumor samples (red dots) and normal tissue samples (teal dots) form two distinct, non-overlapping groups in the two-dimensional space. The first principal component (PC1) accounted for 34.1% of the total variance in the data, while the second principal component (PC2) accounted for 13.7%.

Together, they capture nearly half of the total variation, indicating that the dysregulation of these C-MYC-related genes is indeed the main source of biological difference between the gastric cancer and normal tissue samples in this cohort.

### Diagnostic Potential of Individual Genes

To quantify the potential of each individual gene as a biomarker for gastric cancer diagnosis, Receiver Operating Characteristic (ROC) curve analysis was performed **(Figure 3)**.

**Figure 3:**
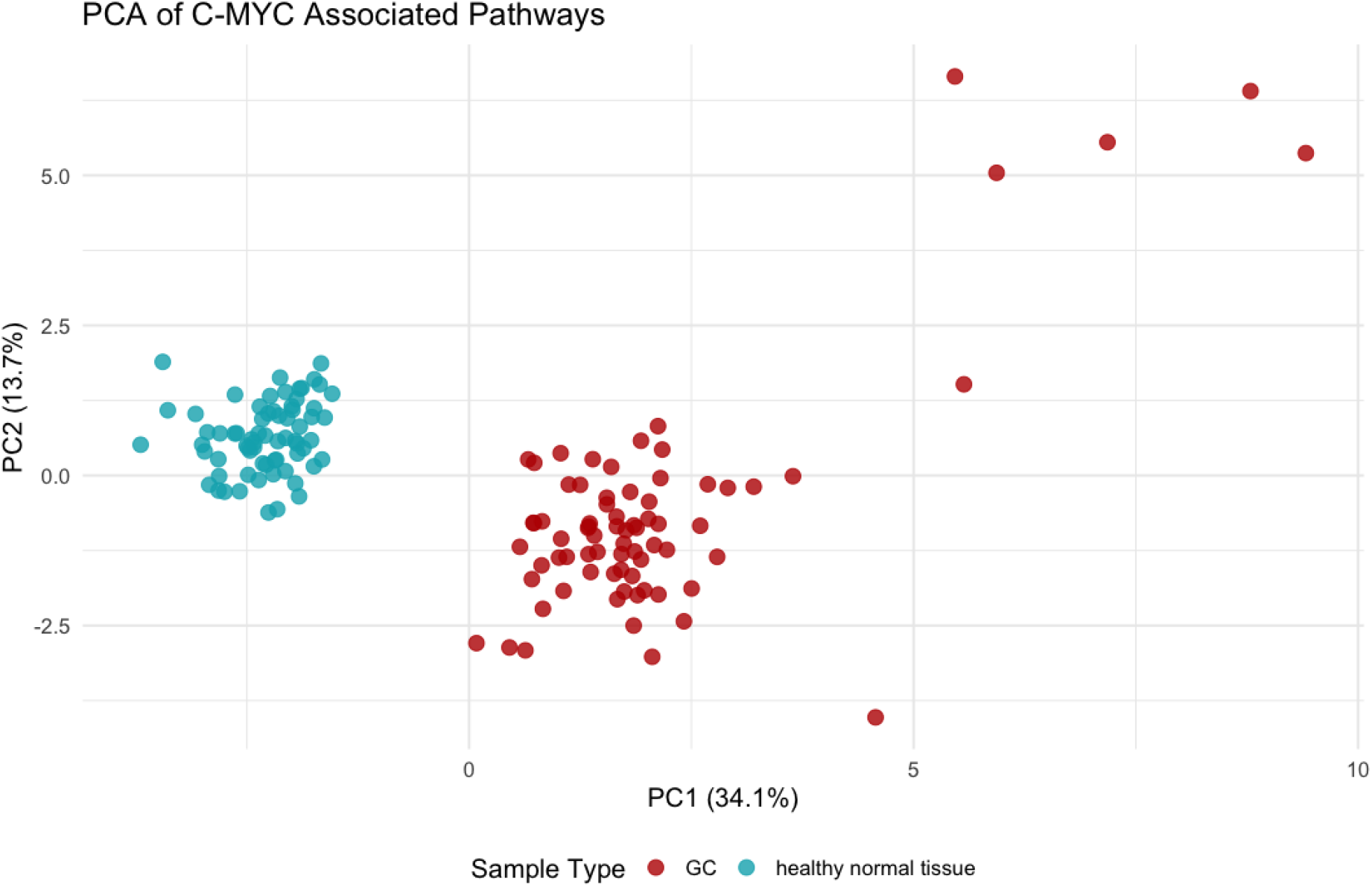
PCA of Pathways Associated with C-MYC: Principal Component Analysis (PCA) of gastric cancer (GC) samples and healthy normal tissues.

**Figure 4:**
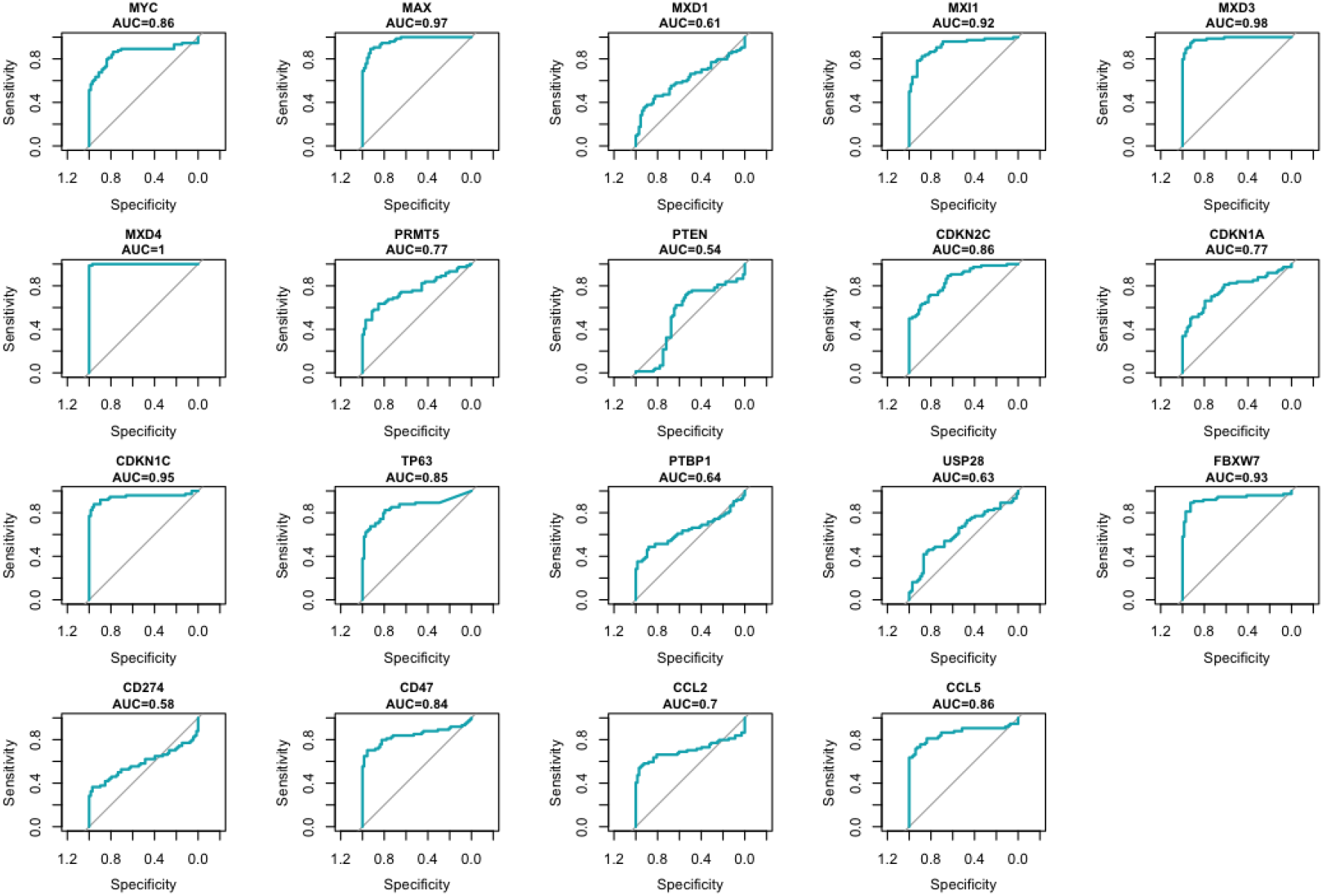
ROC Curve Analysis: Receiver Operating Characteristic (ROC) curve analysis for each of the 21 genes associated with C-MYC.

This approach evaluates the sensitivity and specificity of a marker, a concept also explored in other diagnostic methodologies for MYC status, such as CISH and IHC. The results revealed that several genes possess exceptional diagnostic potential. Notably, the repressor MXD4 demonstrated perfect performance, with an Area Under the Curve (AUC) of 1.00, indicating that its expression could perfectly discriminate between tumor and normal tissues in this cohort. Other genes with near-perfect performance included MXD3 (AUC=0.98) and MAX (AUC=0.97). Additional genes with high diagnostic accuracy (AUC > 0.90) were CDKN1C (AUC=0.95), FBXW7(AUC=0.93), and MXI1 (AUC=0.92). It is noteworthy that MYC itself, despite being the central oncogene, presented a good diagnostic value (AUC=0.86) but was outperformed by several of its partners and antagonists, suggesting that the loss of the regulatory machinery may be an even more sensitive marker for the presence of malignancy.

### Gene Expression by Laurén Histological Subtype

Investigating the clinical relevance of this signature within the tumor cohort, stratification by Laurén classification revealed distinct and statistically significant expression patterns **(Figure 5)**. The expression of MYC was significantly higher in the intestinal subtype compared to the diffuse subtype (p=0.0077). This finding strongly corroborates previous reports in the literature, where C-MYC protein expression and gene amplification at the 8q24.21 locus are more frequently observed in intestinal-type tumors. Supporting this observation, the MYC stability regulator PTBP1 was also significantly more highly expressed in the intestinal subtype (p=0.045). In a striking contrast, an inverse expression pattern was observed for the MYC antagonists. The repressors MXD3 (p=0.00042) and MXI1 (p=0.0024) were significantly more highly expressed in the diffuse subtype. This opposing and divergent signature between the subtypes strongly supports the hypothesis that intestinal and diffuse tumors follow distinct genetic and epigenetic pathways, as suggested by Souza et al. (2013), who pointed to different alteration mechanisms, such as MYC hypomethylation, being more common in diffuse-type tumors.

**Figure 5:**
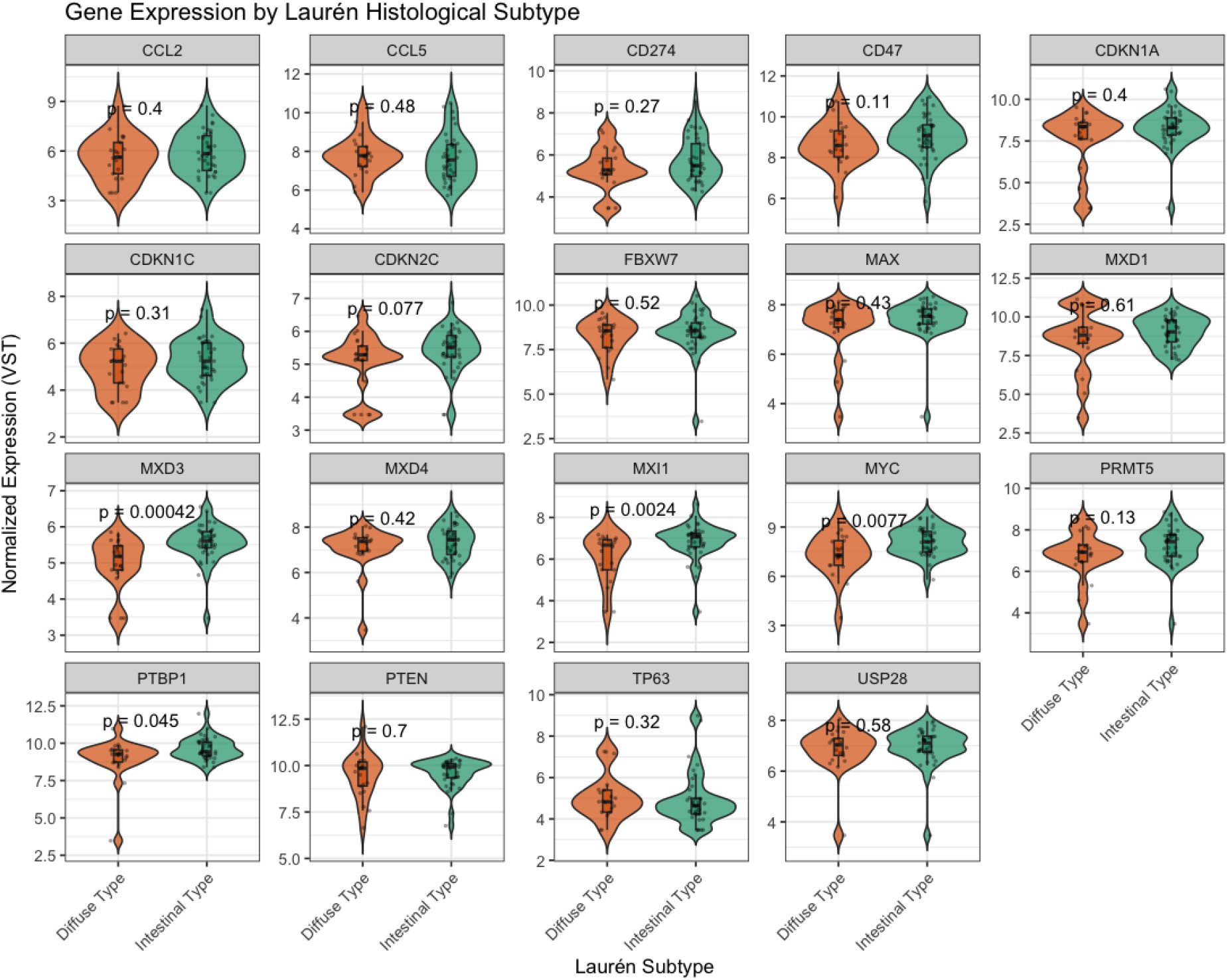
Gene Expression by Laurén Histological Subtype: Violin plots comparing the expression of C-MYC– associated genes between the Diffuse and Intestinal histological subtypes.

### Gene Expression by Neoadjuvant Treatment Status

Finally, the expression of the genes in the signature was evaluated in relation to the patients’ neoadjuvant treatment status **(Figure 6)**. The comparative analysis between tumors from patients who received neoadjuvant therapy (“Sim”) and those who did not (“Não”) revealed no statistically significant difference for any of the 21 genes analyzed. The chemokine gene CCL5 showed a borderline trend (p=0.05) but did not reach the conventional threshold for significance. All other genes, including MYC, showed no association with neoadjuvant treatment status, suggesting that, in this cohort, the baseline expression profile of these genes may not be a primary predictor for the indication of this type of therapy.

**Figure 6:**
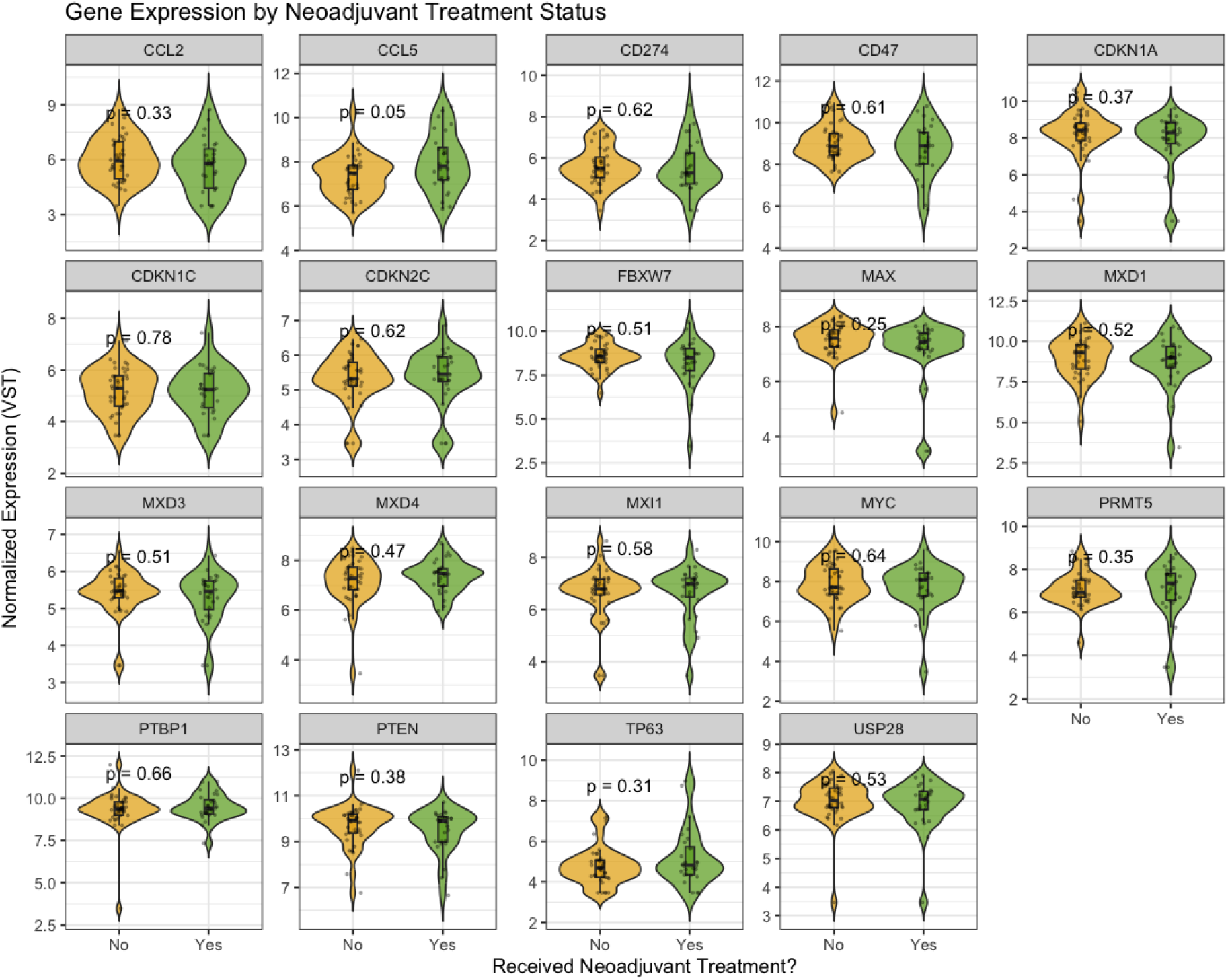
Gene Expression by Neoadjuvant Treatment Status: Violin plots comparing the expression of C-MYC–associated genes based on neoadjuvant treatment status.

## Discussion

This study aimed to perform an integrated bioinformatic analysis of the C-MYC oncogenic network in gastric adenocarcinoma (GAC) to elucidate its expression profile, evaluat its biomarker potential, and correlate it with key clinicopathological factors.

While the role of MYC as an oncogene in gastric cancer (GC) is well-established, the integrated expression pattern of its regulatory network and clinical implications, especially concerning Laurén subtypes and treatment status, remains less understood. Our findings revealed molecular footprints of GC based on C-MYC pathway dysregulation, characterize specific-pathway behaviours across Laurén subtypes, and highlight diagnostic possibilities of its transcriptional repressors, identifying key player genes with significant biomarker potentials.

Our integrated analysis revealed a consistent dysregulation of C-MYC pathway, associated and transcriptional repressors in a significant subset of gastric cancer tumors GAC. This is characterized by a dual hit of MYC activation and loss of its key repressor patterns like MXD4 and MXD3. The central finding of our research is that GC is characterized not only by activation of the MYC oncogene (10) itself but also by the concerted downregulation of its key transcriptional repressors and regulatory partners. This reveals that this oncogenic drive is amplified by a parallel loss of the cellular machinery designed to restrain MYC activity.

The most significantly downregulated genes were the Mad family members MXD4 and MXD3, along with MXI1 and their core partner, MAX. These proteins form heterodimers with MAX to compete with MYC for binding at E-box sequences of target genes, thereby actively repressing transcription and acting as critical tumor suppressors (11). This contributes to a double hit mechanism on the normal regulatory machinery of cell proliferation, enabling cancer cells to also dismantle the machinery designed to suppress it, promoting unchecked proliferative signals, metabolic reprogramming, and avoidance of apoptosis. The finding that MXD3 and MXD4 are more significantly downregulated than MYC is overexpressed is a crucial insight into the dysregulation of the C-MYC pathway in GC, suggesting that tumorigenesis is also being compounded by the simultaneous dismantling of the cellular mechanisms designed to restrain it (12). This model explains why, in our study, the practical absence of repressors like MXD4 (AUC=1.00) serves as a predominant classifier of malignancy than the presence of MYC (AUC=0.86). This suggests that the loss of MXD3/MXD4 may be a more specific diagnostic and prognostic marker for distinguishing malignant tissue from normal tissue. We propose that a diagnostic panel incorporating MXD4 or MXD3 could potentially improve early cancer detection or diagnostic confirmation in biopsies.

However, the downregulation of cell cycle inhibitors like CDKN1C (p57) and CDKN1A (p21) concords to the direct reflection of well-established dual function of MYC pathway as a transcriptional activator and a repressor (13). This coordinated pattern - the upregulation and downregulation of C-MYC pathway, vividly captured by the hierarchical clustering and Principal Component Analysis (PCA), presents a unified and unique identification of a rewired oncogenic network in gastric cancer.

Substantial evidence indicates that while MYC amplification or overexpression occurs in a subset of tumors, the epigenetic silencing or deletion of tumor suppressors like the Mad genes is a widespread phenomenon in cancer (6). However, our results suggest that assessing the expression of these repressors could provide a highly accurate molecular tool for distinguishing malignant from normal gastric tissue thereby potentially surpassing current methods.

Our stratification by Laurén classification provides compelling evidence that intestinal and diffuse subtypes not only differ morphologically but also in their engagement of the C-MYC pathway. The significant overexpression of MYC and its stability regulator PTBP1 in the intestinal subtype is highly consistent with the known pathogenesis of this variant. Intestinal-type GAC often occurs through a stepwise progression commonly known as the Pelayo Correa cascade and is associated with specific genomic instability, including amplifications at the 8q24 locus where MYC is located (14). Our results confirm that MYC-driven oncogenesis is a hallmark of this pathway.

Conversely, this analysis identifies the significant upregulation of the repressors MXD3 and MXI1 in the diffuse subtype as an intriguing counterpoint. This suggests that the nature of diffuse-type GAC may be driven by different underlying mechanisms, and the MYC pathway playing a differently regulated role. This analysis aligns with the findings of Souza et al. (2013), who noted that while MYC amplification was rare in diffuse-type, hypomethylation of the MYC promoter was more common, suggesting a different mode of regulation(10).

Furthermore, the higher expression of Mad proteins in diffuse tumors could be as a result of a failed compensatory mechanism by the cell to counteract other oncogenic drivers, making the drivers in these tumors rely less on gene amplification and more on post-transcriptional and epigenetics mechanisms. This has the potential to trigger an unsuccessful response from repressors like MXD3. However, the role of MXD3 can be context-dependent. MXD3 has been shown to be highly expressed in proliferating cells (15). Furthermore, this suggests that the oncogenic drive in these tumors may rely less on gene amplification and more on post-translational and epigenetic mechanisms, which has the potential to trigger an unsuccessful response from repressors like MXD3.

Following the divergent pathways by the Laurén classification, our results reinforce this biological dichotomy. The intestinal subtype is known to be driven by a classical MYC overexpression mechanism, potentially through gene amplification at the 8q24.21 locus. Conversely, our research identifies the presence of a different underlying mechanism in the diffuse subtype, showing significant upregulation of MXD3 and MXI1. This suggests an alteration in the regulation of the C-MYC pathway, involving mechanisms like the promoter hypomethylation which is associated with the diffuse subtype (10). This fundamental difference in MYC pathway could contribute to the distinct clinical behavior and therapeutic challenges posed by diffuse gastric cancers.

This contrast reinforces the concept that intestinal and diffuse gastric cancers are molecularly distinct diseases, a classification supported by The Cancer Genome Atlas (TCGA) network, 2014, and implies that therapeutic strategies targeting the MYC pathway may be more effective for the intestinal subtype (16).

Interestingly, our analysis found no significant association between the baseline expression of any gene and neoadjuvant treatment status. The relevance of this is that the pre-therapy expression of this specific C-MYC network is not a primary determinant of response to conventional neoadjuvant chemotherapy in this cohort.

We were able to assess the baseline expression; hence it is possible that the MYC expression during treatment or in cellular populations like cancer stem cells could be highly relevant. Future studies with paired pre- and post-treatment samples are needed to explore this possibility. Moreover, the cohort size, particularly for subgroup analysis like treatment status could be more to gain more significance to detect more subtle associations. Also, our analysis uses transcriptomic data while revealing crucial expression patterns, which offers a limited overview of the overall C-MYC pathway dysregulation. Future research should integrate other omics data, including proteomics to evaluate protein expression and activation status, and epigenomics in order to gain a more comprehensive understanding of the molecular alterations driving gastric adenocarcinoma.

Our integrated analysis using transcriptomic data reveals that gastric adenocarcinoma is defined by a coordinated dysregulation of the C-MYC oncogenic network, showing both the activation of the oncogene and the silencing of its key repressors. It also offers a deep insight into C-MYC pathway dysregulation in gastric cancer and identifies significant molecular signatures that provide a powerful tool for diagnosis, with the loss of repressors like MXD4 being a remarkably sensitive marker. This highlights the potential of C-MYC antagonists as prognostic and diagnostic biomarkers, paving the way for more personalized therapeutic approaches.

## Acknowledgment

The authors would like to thank the Oncology Research Center, the Human and Medical Genetics Laboratory, and the Anatomical Pathology Laboratory at João de Barros Barreto University Hospital (HUJBB – UFPA) for their invaluable technical and laboratory support. Our gratitude also goes to the High-Performance Computing Center (CCAD) at the Federal University of Pará for access to the Apollo 2000 cluster, which was crucial for our analyses.

## Data availability statement

The original contributions presented in the study are included in the article. Further inquiries can be directed to the corresponding author.

